# Can BioSAXS Detect Ultrastructural Changes Of Antifungal Compounds In *Candida Albicans*? – An Exploratory Study

**DOI:** 10.1101/2023.05.12.540542

**Authors:** Kai Hilpert, Christoph Rumancev, Jurnorain Gani, Dominic W. P. Collis, Paula Matilde Lopez-Perez, Vasil M. Garamus, Ralf Mikut, Axel Rosenhahn

## Abstract

The opportunistic yeast *Candida albicans* is the most common cause of candidiasis. With only four classes of antifungal drugs on the market, resistance is becoming a problem in the treatment of fungal infections, especially in immunocompromised patients. The development of novel antifungal drugs with different modes of action is urgent. In 2016, we developed a groundbreaking new medium-throughput method to distinguish the effects of antibacterial agents. Using small-angle X-ray scattering for biological samples (BioSAXS), it is now possible to screen hundreds of new antibacterial compounds and select those with the highest probability for a novel mode of action. However, yeast (eukaryotic) cells are highly structured compared to bacteria and the action of an antifungal drug might leave most structures unchanged. In the pioneering work described here, we explored the possibility if BioSAXS can be used to measure the ultrastructural changes of *Candida albicans* directly or indirectly induced by antifungal compounds. For this exploratory study, we used the well-characterized antifungal drug flucytosine. BioSAXS measurements were performed on the synchrotron P12 BioSAXS beamline, EMBL (DESY, Hamburg) on treated and untreated yeast *C. albicans*. BioSAXS curves were analysed using principal component analysis (PCA). The PCA showed that flucytosine-treated and untreated yeast were separated. Based on that success further measurements were performed on five antifungal peptides (1. Cecropin A-melittin hybrid [CA(1-7)M(2-9)], KWKLFKKIGAVLKVL; 2. Lasioglossin LL-III, VNWKKILGKIIKVVK; 3. Mastoparan M, INLKAIAALAKKLL; 4. Bmkn2, FIGAIARLLSKIFGKR; and 5. optP7, KRRVRWIIW). The ultrastructural changes of *C. albicans* indicate that the peptides may have different modes of action compared to Flucytosine as well as to each other, except for the Cecropin A-melittin hybrid [CA(1-7)M(2-9)] and optP7, showing very similar effects on *Candida albicans*. This very first study demonstrates that BioSAXS shows great promise to be used for antifungal drug development. The use of such a tool like BioSAXS could accelerate and de-risk antifungal drug development, however, further experiments are necessary to establish this application.

## 1 Introduction

Similar to the problem of antimicrobial resistance in bacteria, human pathogenic yeast and fungi also show an increase in resistance (Bhattacharya et al., 2020; Kean and Ramage, 2019; Murray et al., 2022). With the growing numbers of immunocompromised patients, the resistance of opportunistic fungi to antifungal drugs is of concern since only four classes of antifungals are available for treatment. For example, patients who are undergoing cancer treatment (chemotherapy), have transplants, have diabetes, or are infected by the human deficiency virus (HIV-1/2) are at higher risk of getting infected by fungi, especially yeast. In addition, immunocompetent individuals can also experience symptoms of yeast overgrowth, for example, topical, oral, vulvovaginal, and intra-abdominal, causing estimated annual cases of more than 130 million (Arastehfar et al., 2019). The most common cause of candidiasis is the opportunistic yeast *Candida albicans*.

Small-angle X-ray scattering (SAXS) is a powerful tool to study ultrastructures and their changes in macromolecules and nanomaterials in solution or suspension (Boldon et al., 2015; Byer et al., 2023; Honecker et al., 2022; Sartori and Marmiroli, 2022; Sun et al., 2022). The combination of a short wavelength and a large penetration depth of the photons used allows the investigation of large volumes that contain millions of objects in less than a second, providing unique analytical possibilities (Franke and Svergun, 2020). That allows for gathering structural information on disordered systems and, due to the large penetration depth, resolving the inner structures of such systems. After the sample has been irradiated, the scattered photons are recorded with a spatially resolved 2D detector.

In our pioneering work over the last years, we have shown that the ultrastructural changes caused by the treatment of bacteria with antibiotics can be detected by BioSAXS (Hilpert et al., 2021; Rumancev et al., 2022; Von Gundlach et al., 2016a, 2016b, 2019). In this approach, BioSAXS is used as a structurally sensitive fingerprint method that reveals subtle differences in the intracellular structure of the pathogens, which are caused by the drug treatment directly or as a response of the microorganism towards the assault (Rumancev et al., 2022). The intracellular structural changes occur on different length scales, and a combination of SAXS and ultra small-angle X-ray scattering (USAXS) allowed to separate contributions from at least four different structure sizes, which were connected to major bacterial properties such as the shape or size of the ribosomes (Von Gundlach et al., 2016b). Recently, it was shown that also neutron scattering can provide additional useful information about the bacterial ultrastructure (Semeraro et al., 2022). BioSAXS can be used for Gram-positive as well as Gram-negative bacteria (Von Gundlach et al., 2019). The differences in the scattering curves could be analysed using a principal component analysis (PCA), and plots of PCA 1 over PCA 2 can be used to visualise the results. We could demonstrate that those distances between data points for untreated versus treated with different antibiotics correlated with their different modes of action (Von Gundlach et al., 2016a). However, it was unclear if BioSAXS could be used to support the development of urgently needed new antifungal drugs. Fungi are true eukaryotes and therefore are immensely more structured than eukaryotes. These various structures will be picked up by BioSAXS, and the question we address in this study is whether ultrastructural changes caused by the antifungal drug can be detected above the background of all existing structures within the cell. The focus of this study was exploratory and fundamental: can BioSAXS be used to discriminate between untreated yeast (*Candida albicans*) cells and cells treated with an antifungal compound?

Flucytosine was used as a well-known antifungal drug. Flucytosine can be deaminated in yeast, but not in mammalian cells, hence its specificity and usage as a drug. The deaminated version will be incorporated into RNA and inhibit protein biosynthesis. In a further enzymatic reaction by uridine monophosphate pyrophosphorylase, 5-fluorodeoxyuridine monophosphate is produced. This compound inhibits thymidylate synthetase, and consequently, DNA synthesis is inhibited, also leading to further disruption in protein synthesis (Vermes et al., 2000). The antifungal compound was incubated with the microorganism for 40 minutes and then chemically fixed. SAXS curves were acquired and analysed by principal component analysis to group similar intracellular morphological responses to the drug treatment.

## Materials and Methods

### Peptides

Antimicrobial peptides were produced by automated solid-phase peptide synthesis (SPPS) on a MultiPep RSI peptide synthesizer (Intavis, Tuebingen; Germany) and purified to the homogeneity of >90% by liquid chromatography-electrospray ionisation mass spectrometry (LC-ESI-MS, Shimadzu LC2020 system, Shimadzu, Milton Keynes, United Kingdom). For more details please refer to (Grimsey et al., 2020).

### Yeast Strain

In this project, we used a clinical isolate of *Candida albicans* obtained from Tim Planche (St. George’s, University of London).

### Bacteriological Media and Culture Conditions

Mueller-Hinton broth (MHb) (Merck, Life Science UK Limited, Dorset, UK)) was prepared to concur with the manufacturer’s directions and supplemented with 2% glucose to grow *C. albicans*. Liquid cultures were incubated at 30°C for 18–20 h on a shaker, and culture on solid media was incubated for 18–24 h at 30°C.

### Minimal Inhibitory Concentration Determination

When it comes to determining the antimicrobial activity of antimicrobial peptides (AMPs) there are unfortunately many different approaches reported. Many AMPs are negatively affected by positively charged ions and other components of growth media, therefore Mueller-Hinton broth (MHb) is frequently used since it supports the growth of many organisms and seems to interfere little with the activity of most AMPs. The mode of action of Flucytosine and antifungal peptides were determined using a protocol (option E) of the highly cited publication by Wiegand et al. optimized for the work with antimicrobial peptides (Wiegand et al., 2008). This protocol uses MHb (0.2 % beef extract, 1.75 % casein hydrolysate). To adopt that media for the growth of yeast 2% glucose was added. For comparison, YPD complex medium (1% yeast extract, 2% peptone, 2% dextrose) is often used for the growth of yeast. Where this media provides the best activity for antimicrobial peptides the results should be carefully used when comparing MIC values to other antifungals measured in different growth media. A broth microdilution assay was applied using 6 × 10^5^ CFU/ml at 30°C. MICs were determined by the unaided eye, as the concentration in which no growth was observed.

### Sample Preparation, Measurement and Data Evaluation for BioSAXS

A detailed description of the sample preparation for BioSAXS, the BioSAXS measurement and consequently the data evaluation methods can be found in the open-source publication by von Gundlach et al. (Von Gundlach et al., 2019). Briefly, an aliquot of an overnight culture of a clinical isolate of *Candida albicans* was diluted in Mueller-Hinton broth (MHb) and incubated for 2.5 h at 30°C on a shaker (225 rpm) to achieve log phase growth. The *C. albicans* culture was adjusted to have 1.2 × 10^6^ CFU/ml and aliquoted. The aliquots were then mixed with either sterile water or antifungal solution (2 x MIC) to provide 6 × 10^5^ CFU/ml aliquots. Those were further incubated for 40 min at 30°C at 225 rpm. All aliquots were then centrifuged at 10,000 rpm and washed 3 times in ml 0.1 M PIPES (piperazine-N,N′-bis(2-ethanesulfonic acid)) buffer, pH 7.0. After washing, the pellet was resuspended in 1 ml of 2.5% glutaraldehyde v/v in PIPES buffer and fixated for 1 h at room temperature at 225 rpm. Consequently, the samples were washed 3 times with PBS buffer and resuspended in 100 μl of PBS and stored at 5°C.

The SAXS measurements were performed on the P12 BioSAXS beamline, EMBL (DESY, Hamburg), using a double crystal Si (111) monochromator. An energy of 10 keV with a 10^13^ s^-1^ photon flux was used. 30 μl of each sample was injected into the measuring chamber by an automatic sample robot (Arinax, BioSAXS), the focus of the X-ray beam was 0.2 × 0.12 mm. The scattering signal was recorded with a Pilatus 2M detector (Dectris, Switzerland). For each sample, 20 measurements were made with 0.05 s exposure. Before and after each measurement the background was measured and automatically subtracted from the SAXS curves. Data analysis and evaluation were done using the open-source software SciXMiner (“Peptide Extension”) (Mikut, 2010; Mikut et al., 2017). To determine differences in the scattering curves between treated and untreated samples as well as between the different treatments a principal component analysis (PCA) was applied to the scattering curves.

## Results

In this project, our main objective was to test whether BioSAXS can discriminate between untreated *C. albicans* cells and those treated with Flucytosine. The MIC of Flucytosine was determined to be 8 μg/ml. For bacterial culture, we found that 1×10^8^ CFU/ml is optimal for getting enough scattering signal to meaningfully analyse the curves from the BioSAXS experiment (Von Gundlach et al., 2016a). *Candida albicans*, however, is a eukaryote that, in its yeast form is about 5-10μm in diameter, whereas, for example, *Staphylococcus aureus* is only 0.5-1.0 μm in diameter. In addition, eukaryotic cells are more structured and therefore may have a stronger scattering signal in a BioSAXS experiment than prokaryotic cells. Consequently, for the first experiments, we used only 6 × 10^5^ CFU/ml and combined this with 2xMIC for Flucytosine. Flucytosine was incubated for 40 minutes and then the cells were fixated. Fortunately, the scattering curves showed good signal strength and could be used for further analysis. The scattering curves are presented in Figure 1.

**Figure 1.**
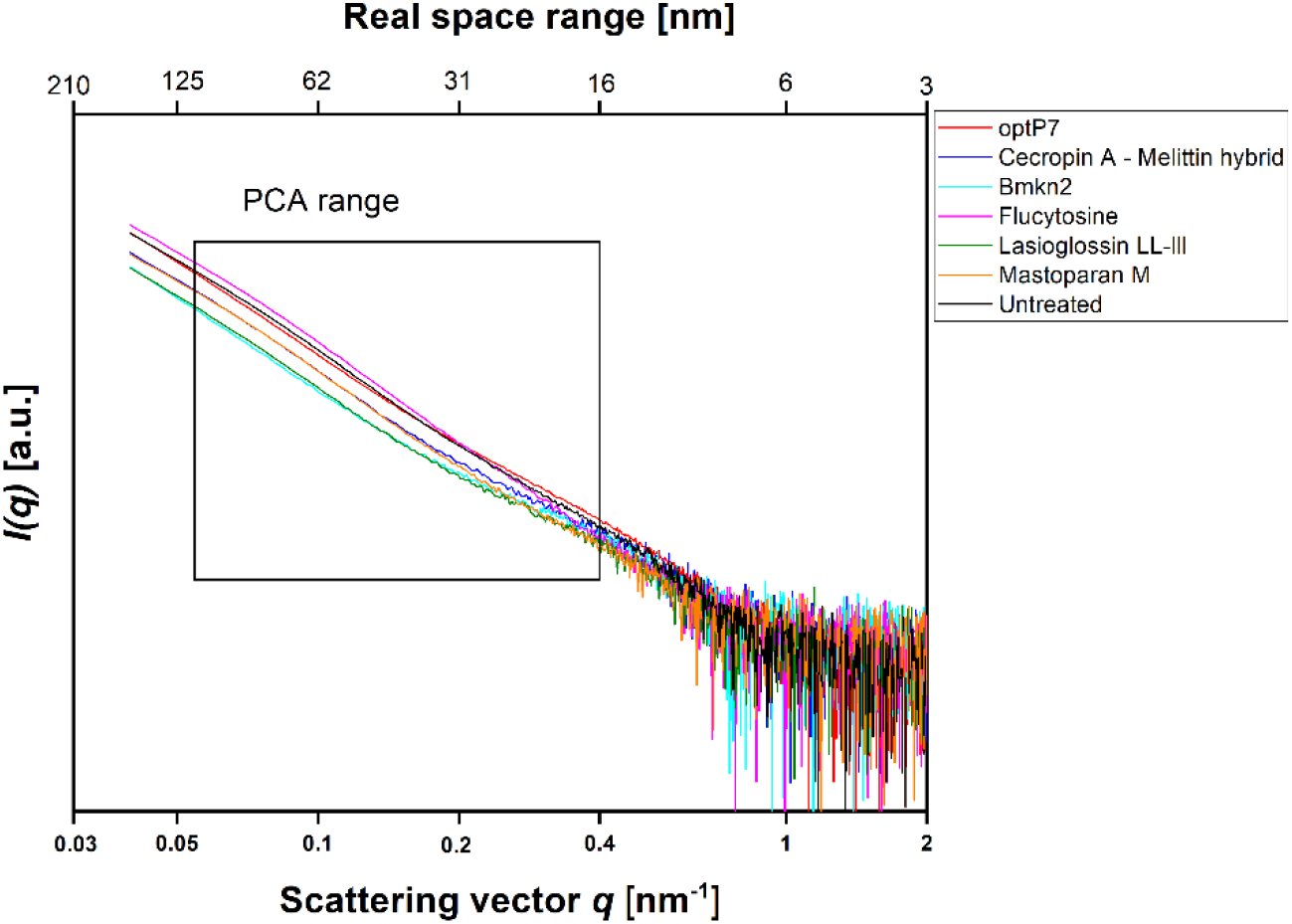
Small angle X-ray scattering data from *Candida albicans* as measured at the P12 BioSAXS beamline at PETRA III (Hamburg, Germany) at a photon energy of 10 keV. Untreated *C. albicans* as well as treated (40 min, 2xMIC) with antifungal peptides and Flucytosine were measured. The box indicates the range that was used to calculate the principal component analysis (PCA).

The data was analysed by open-source data mining MATLAB® Toolbox SciXMiner, using the “Peptide Extension” tool (Mikut, 2010; Mikut et al., 2017). A principal component analysis was performed to extract differences between the curves and for visual presentation, the values of the linear coefficients for the first two principal components were plotted, see Figure 2a.

**Figure 2.**
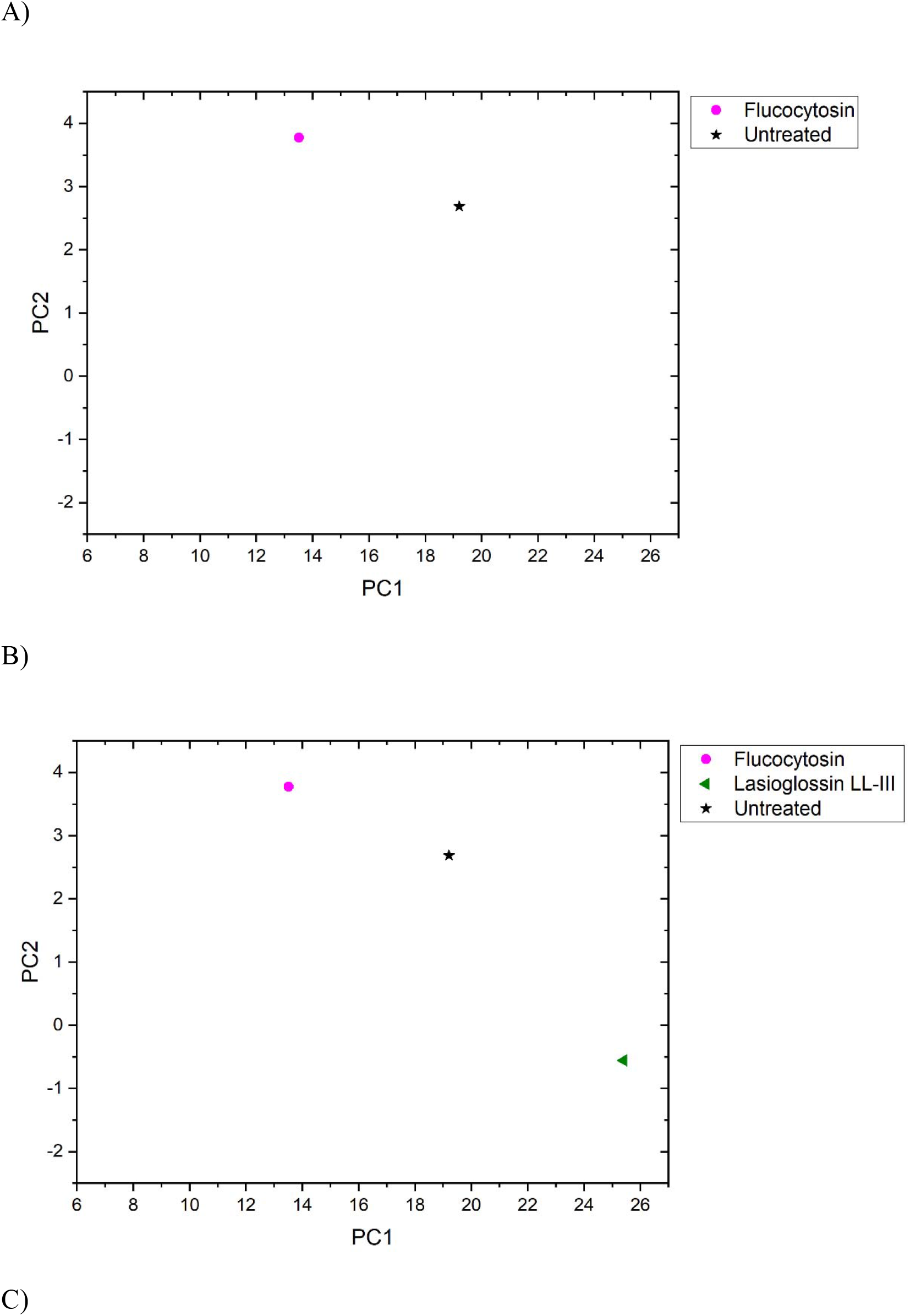
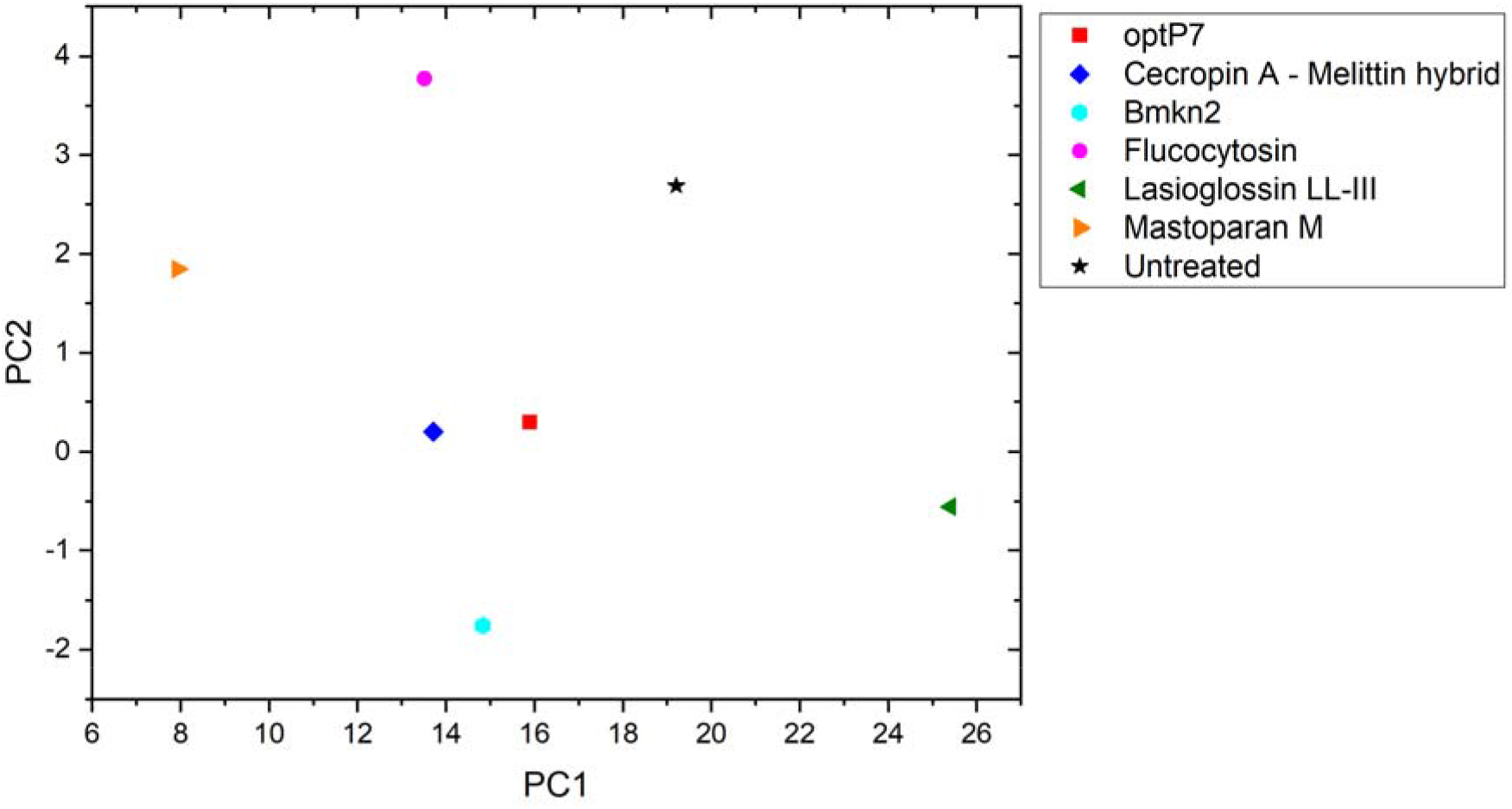
The linear coefficients of the first two principal components (PC) discriminate morphological changes and modes of action. A) Flucytosine and untreated, B) Flucytosine, Lasioglossin LL-III and untreated and C) Flucytosine, Lasioglossin and various antifungal peptides.

The data are measurements of about 100,000 cells that were measured 20 times to obtain the scattering curve. In consequence, the data has high statistical robustness. Flucytosine induced ultrastructural changes after 40 min that separates well from the untreated sample. It shows a reduction of PC1 by about 6 units and an increase of PC2 by about 1 unit. This was the first time demonstrating that BioSAXS can be used to investigate antifungal compounds directly in yeast. That opens the way for further systematic studies similar to those done for antibacterial compounds on different bacteria. The main objective of this study was successfully fulfilled. Based on that success, a few antifungal peptides were selected for a further experiment.

Antimicrobial peptides are a class of compounds found in all four kingdoms of life that act against bacteria, fungi, viruses, and parasites. There are various databases for antimicrobial peptides, for example, APD, CAMP and DRAMP and DBAASP (Gawde et al., 2023; Kang et al., 2019; Pirtskhalava et al., 2021; Wang et al., 2016). DBAASP for example, currently lists 5595 antifungal peptides (Pirtskhalava et al., 2021). There is a huge potential to develop novel antifungals using these peptides as templates. Peptide libraries on cellulose can be used to investigate this large pool of compounds economically (Ashby et al., 2014, 2017; Knappe et al., 2016; López-Pérez et al., 2017; Mania et al., 2010). Of special interest are peptides that demonstrate a different mode of action than conventional antifungal drugs. Unfortunately, screening even small numbers of peptides to determine their mode of action is an enormous task and a great financial burden. Therefore, we decided to test one antifungal peptide (Lasioglossin LL-III) in a BioSAXS experiment. Lasioglossin LL-III was isolated from the venom of the eusocial bee, *Lasioglossum laticeps* (C□er□ovský et al., 2009). The peptide displays favourable features for further drug development, including strong antimicrobial activity combined with low hemolytic and cytotoxic activity. The peptide can interact with charged membranes in a non-disruptive manner, depolarizes the membrane, and binds to DNA, suggesting DNA as an intracellular target (Battista et al., 2021). The peptide was able to kill various yeasts, including *C. albicans*, within 15 minutes and was shown to depolarize the plasma membrane (Kodedová and Sychrová, 2016). The antifungal activity of Lasioglossin LL-III against *C. albicans* was determined at 11 μM using a low inoculum of 1.2–7.5×10^3^ CFU per well (Slaninová et al., 2011). In addition, it was shown that the peptide inhibited temperature-induced morphotype changes toward hyphae and pseudohyphae (Vrablikova et al., 2017). The authors also showed that the peptide is an inhibitory agent in the DBA/2 murine model of vulvovaginal candidiasis and therefore a potential new drug candidate. The minimal inhibitory concentration (MIC) was determined using 6 × 10^5^ CFU/ml at 30°C to be 36 μM (Table 1). At two times the MIC and an incubation time of 40 min, *C. albicans* samples were prepared and measured by BioSAXS. The results are presented in Figure 2b. Surprisingly, there was a strong separation between the effects of Flucocytosin and Lasioglossin LL-III. Whereas Flucytosine acts on protein synthesis by RNA modification and also on DNA synthesis, Lasioglossin LL-III permeabilizes the membrane and interacts rather unspecifically with all DNA in the cell. BioSAXS measurements show that the ultrastructural changes are different. Especially PC1 is increased by 6 units compared to the untreated cells. In consequence, Flucytosine and Lasioglossin LL-III are separated by 12 units at the PC1 level. The PC2 value for the peptide is decreased compared to the untreated cells, whereas for Flucytosine it increased.

**Table 1.**
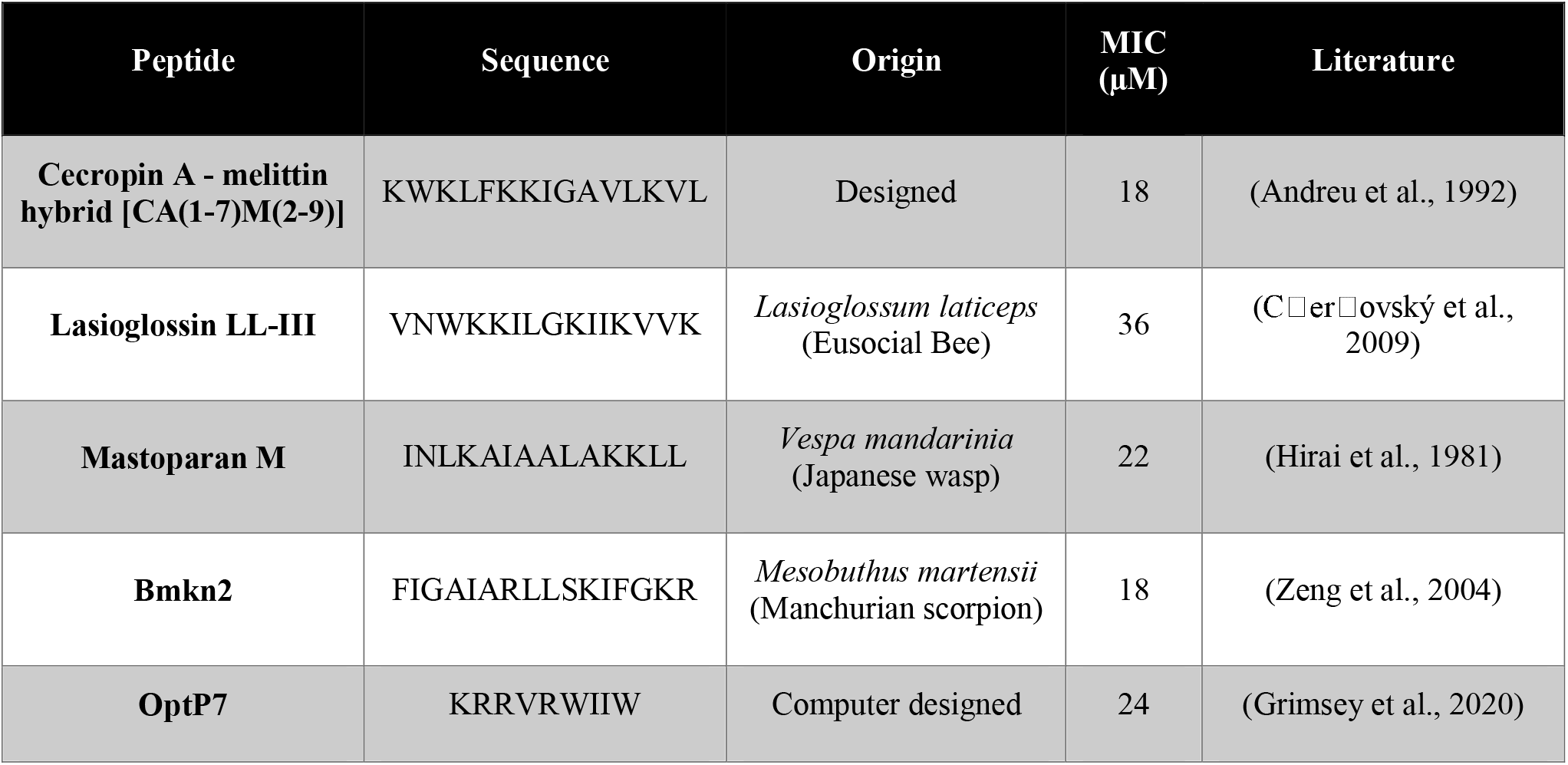
Minimum inhibitory concentrations (MIC) in μM. Data are for n = 3 with modal values reported; all values were determined in Mueller-Hinton broth (MHb) supplemented with 2% glucose using 6 × 10^5^ CFU/ml *C. albicans*. All peptides are C-terminally amidated.

As data obtained on the well-described Flucytosine and Lasioglossin was very encouraging, four less-studied peptides were analyzed and *C. albicans* cells treated with these compounds were measured by BioSAXS. The 16-mer peptide Bmkn2 was identified in the scorpion venom of *Buthus martensii Karsch* (Zeng et al., 2004). The peptide displays strong activity against gram-positive and gram-negative bacteria (Almaaytah and Albalas, 2014). Mastoparan M is a 14-mer cationic peptide that was isolated from the venom of the wasp *Vespa mandarinia* (Hirai et al., 1981). It is active against gram-positive and gram-negative bacteria and shows very little hemolytic activity. In addition, it shows pro-inflammatory activity as well as anticancer activity (Wu et al., 1999; Wu and Li, 1999). The hybrid peptide from Cecropin A from *Hyalophora cecropia* (giant silk moth) and Melittin from Apis mellifera (European honey bee), called CA(1-7)M(2-9) has strong antibacterial activity against gram-positive and gram-negative bacteria, follow-up research shows that it is also active against anaerobe species (Andreu et al., 1992; Edlund et al., 1998; Hultmark et al., 1980; Raghuraman and Chattopadhyay, 2007). The peptide optP7 was obtained from a previously reported peptide library containing 3,000 members to better understand short antimicrobial peptides (Mikut et al., 2016). Based on the analysis of this data we developed a novel prediction method (unpublished results) for AMPs with low hemolytic activity. One selected candidate with strong antimicrobial activity and low hemolytic activity was the peptide optP7. The peptide optP7 was selected for studying lipidation and glycosylation (Grimsey et al., 2020), cyclisation, and grafting into a cyclotide (Koehbach et al., 2021) that could be further optimized by procedures described previously (Hilpert et al., 2005, 2006; Mardirossian et al., 2018, 2019, 2020). In addition, the peptide was used to create hybrid peptides with improved activity and BioSAXS analysis showed different ultrastructural changes compared to both parental peptides (Hilpert et al., 2021). The MICs of all peptides are given in Table 1.

Flucytosine induced more distinct different ultrastructural changes in *C. albicans* than all other antifungal peptides measured (see Figure 2c). It shows the highest PC2 value in the set and has the second-lowest PC1 value. Its specific way of interfering with RNA, DNA, and protein synthesis leads to changes in the cells that do not occur with the treatment of the yeast with the five peptides tested. The highest PC-1 value and the second-lowest PC-2-value are seen in Lasioglossin LL-III. The peptide Lasioglossin LL-III from the eusocial bee is therefore also well separated from all other antifungal compounds tested. Its ability to bind unspecifically to DNA and interfere with replication and RNA synthesis seems to induce rather unique ultrastructural changes. The lowest PC2 value was observed with the peptide Bmkn2 from scorpion venom. The lowest PC1 value was observed with the peptide Mastoparan M from the Japanese wasp. The 15-mer peptide Cecropin A-melittin hybrid [CA(1-7)M(2-9)] and the 9-mer peptide optP7 show very similar ultrastructural changes after 40 min of treatment. In contrast to the other peptides, they are both designed or computer-predicted. Short AMPs like optP7 have been shown to bind to ATP and inhibit ATP-dependent processes (Hilpert et al., 2010) and, in addition, depolarize bacterial membranes (Hilpert et al., 2006). Comparing *C. albicans* treated with all antifungal compounds versus the untreated sample, the differences between PC1 (ΔPC1) and PC2 (ΔPC2) can be calculated. Only Flucytosine shows a positive ΔPC1, demonstrating the different mechanism in comparison to the peptides. Interestingly, only Lasioglossin LL-III has a positive ΔPC1. In all other cases ΔPC1 and ΔPC2 are negative.

## Discussion

The development of resistance against antifungal drugs threatens successful treatment. Novel antifungal substances with new modes of action are urgently needed. Determining the mode of action is still time- and resource-consuming, so screening is not possible. Because of our pioneering work on BioSAXS on bacteria, we demonstrated that screening for novel modes of action is now possible. However, prokaryotes are not very structured in comparison to eukaryotes. In light of the presence of additional structures in eukaryotes, like the cell core, Golgi apparatus, mitochondria, and endoplasmic reticulum, it seemed less likely that BioSAXS could detect changes since so many structures of the yeast cells most likely stay intact when treated with antifungals. We wanted to investigate whether it is possible to detect ultrastructural changes in an eukaryote. Surprisingly, such ultrastructural changes could indeed be measured by BioSAXS using Flucytosine as an antifungal drug. To the best of our knowledge, this is the first study to show that BioSAXS can be used successfully to differentiate between untreated *C. albicans* cells and those treated with an antifungal compound. Based on this success, different antifungal peptides were used and all of them showed differentiating effects from the untreated cells. This is the first proof-of-principle study with an exciting outcome, however, this is the very beginning of the work, and various studies need to be performed to verify the data, expand on known antifungal drugs, and perform microscopy in parallel. These experiences are important for learning more about the potential applications, robustness, and limitations of this method. The final aim is to use BioSAXS as a method to develop novel antifungal drugs with new modes of action faster, cheaper and with less risk involved, see Figure 3.

**Figure 3.**
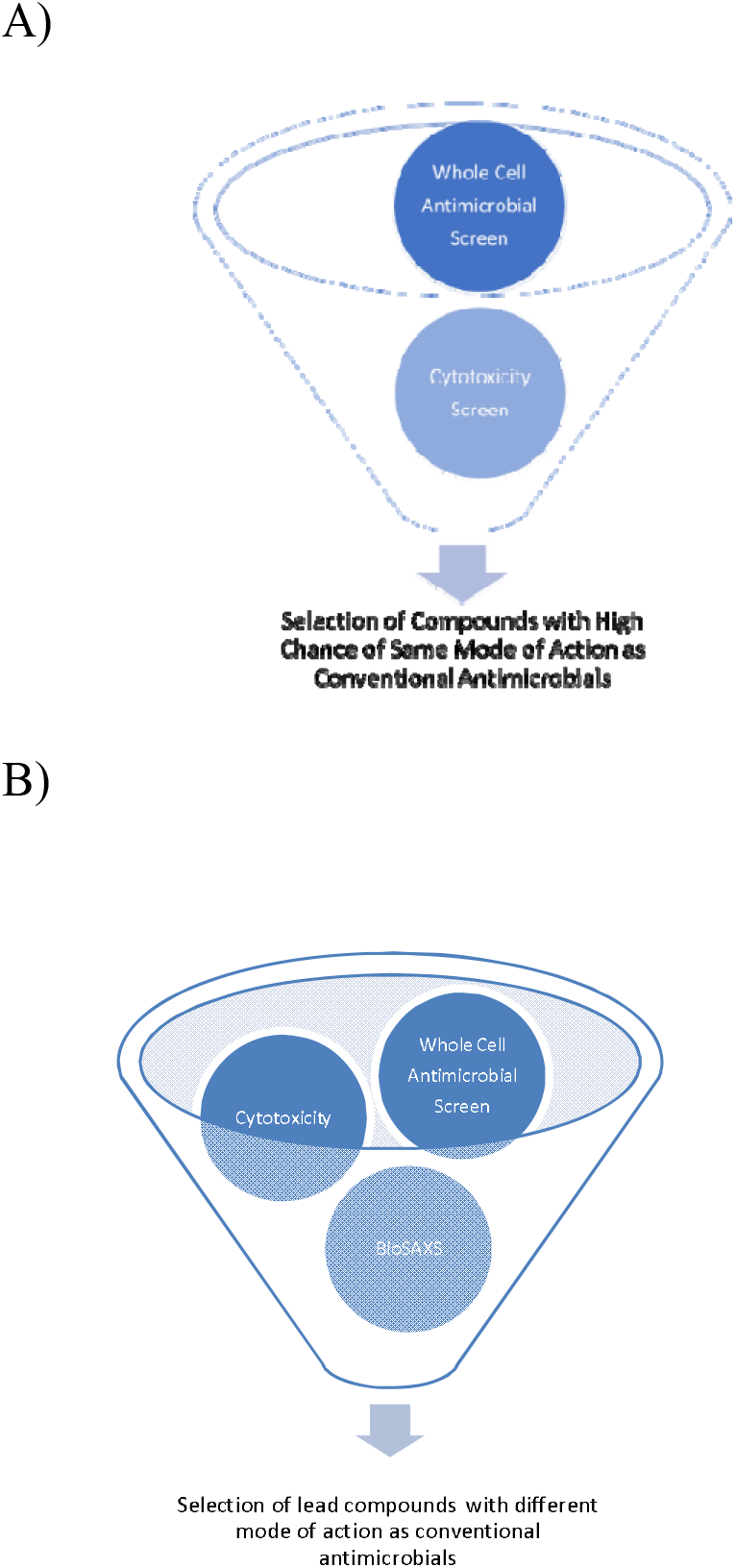
Schematic representation of early drug discovery process using a large compounds library against a targeted microorganism by a whole-cell screen, without BioSAXS, A) and with BioSAXS, B). Taken from (Rumancev et al., 2022)

## Author Contributions

KH conceptualization, resources (peptides, assay development), data analysis, supervision, funding acquisition, writing—original draft preparation, writing—review and editing; CR investigation, data curation (BioSAXS, data analysis and presentation); JG investigation, data curation (peptide synthesis and purification, MICs, BioSAXS sample preparation); PML-P; investigation, data curation (peptide synthesis and purification), supervision; DWPC; investigation, data curation (peptide synthesis and purification), supervision, VG resources (BioSAXS); RM resource (BioSAXS data analysis), funding acquisition; AR conceptualization, resources (BioSAXS and data analysis), data analysis and discussion, supervision, funding acquisition, writing—review and editing. All authors have read and agreed to the published version of the manuscript.

## Acknowledgements

We acknowledge the team of P12 BioSAXS beamline at EMBL, Hamburg for excellent support. KH thanks the Universe/life for the opportunity to continue on and despite the odds to be able to keep researching. Funding by the BMBF project 05K22PC1 is kindly acknowledged.

